# Leucine Rich Repeat Kinase 2 is not required for lysozyme expression in intestinal Paneth cells

**DOI:** 10.1101/2024.03.07.582590

**Authors:** Anna Tasegian, Dina Dikovskaya, Molly Scott, Tom Helps, Tosca Meus, Mairi H. McLean, Mahima Swamy

## Abstract

Genetic variants in Leucine-rich Repeat Kinase 2 (*LRRK2)* gene have been linked to Parkinson’s disease (PD) and are also associated with inflammatory bowel diseases (IBD), specifically Crohn’s disease (CD), a transmural inflammation that can affect the entire length of the gastrointestinal tract and is commonly seen in the ileum^1^. In ileal CD, defects in specialized intestinal epithelial cells known as Paneth cells are believed to drive disease pathogenesis^2,3^. Paneth cells contribute to mucosal defense by secreting antimicrobial peptides including lysozyme and to the maintenance of intestinal stem cells by secreting growth factors. A previous article published by Zhang *et al*^4^ in *Nature Immunology* identified a key role for LRRK2 in selective sorting and secretion of lysozyme in Paneth cells. However, after extended analyses, we find that LRRK2 is not required for lysozyme expression in the murine gut and is not expressed in either murine or human Paneth cells.

## Main Text

LRRK2 is a large multi-domain protein kinase that has been genetically implicated in multiple diseases, most notably Parkinson’s disease (PD), but also in immune-mediated diseases such as Crohn’s disease (CD)^1^. Due to accumulating evidence of abnormal Paneth cells based on the morphology of lysozyme-containing granules in ileal sections from CD patients^2,3,5^, there has been considerable interest in a study published in 2015 showing a role for LRRK2 in lysozyme sorting into granules in Paneth cells^4^. In their study^4^, Zhang *et al* found that *Lrrk2*^*-/-*^ (LRRK2 KO) mice were more prone to intestinal infection. As LRRK2 deficiency was not associated with changes in the gut microbiota, they then evaluated the bactericidal activity of Paneth cells. The lack of lysozyme-positive staining in ileum sections of LRRK2 KO mice, and the defective bactericidal activity of supernatants and cell lysates from crypts and organoids isolated from LRRK2 KO mice, led the authors to argue that genetic ablation of LRRK2 led to lysozyme deficiency in Paneth cells. They went on to show that LRRK2 within Paneth cells acted downstream of intestinal microbes and the pathogen-recognition receptor NOD2 to mediate the correct sorting of lysozyme to secretory granules, and in the absence of LRRK2 lysozyme was instead mistargeted to lysosomes for degradation.

The findings of the study by Zhang et al^4^ strongly implicated LRRK2 in the ability of Paneth cells to produce lysozyme and suggested an important mechanistic link between LRRK2 mutations and Crohn’s disease, which deserved further exploration. Surprisingly, in our analyses of lysozyme expression in Paneth cells of LRRK2 KO^6^ mice (**Fig. 1a**) and in LRRK2 R1441C mice (Parkinson’s disease associated mutation, data not shown), we did not find any defects in lysozyme expression or sorting into granules. Notably, a thorough quantitative analysis of multiple ileal crypts (85-112 crypts from 8 random images per mouse) from LRRK2 WT and LRRK2 KO (7 mice per genotype) mice revealed an equivalent total lysozyme staining intensity in LRRK2 WT and LRRK2 KO mice (**Fig. 1a**). However, we did note a slight reduction in the maximum lysozyme intensity/crypt in the LRRK2 KO mice, which suggests that the lysozyme distribution in LRRK2 KO mice is more diffuse.

**Figure 1.**
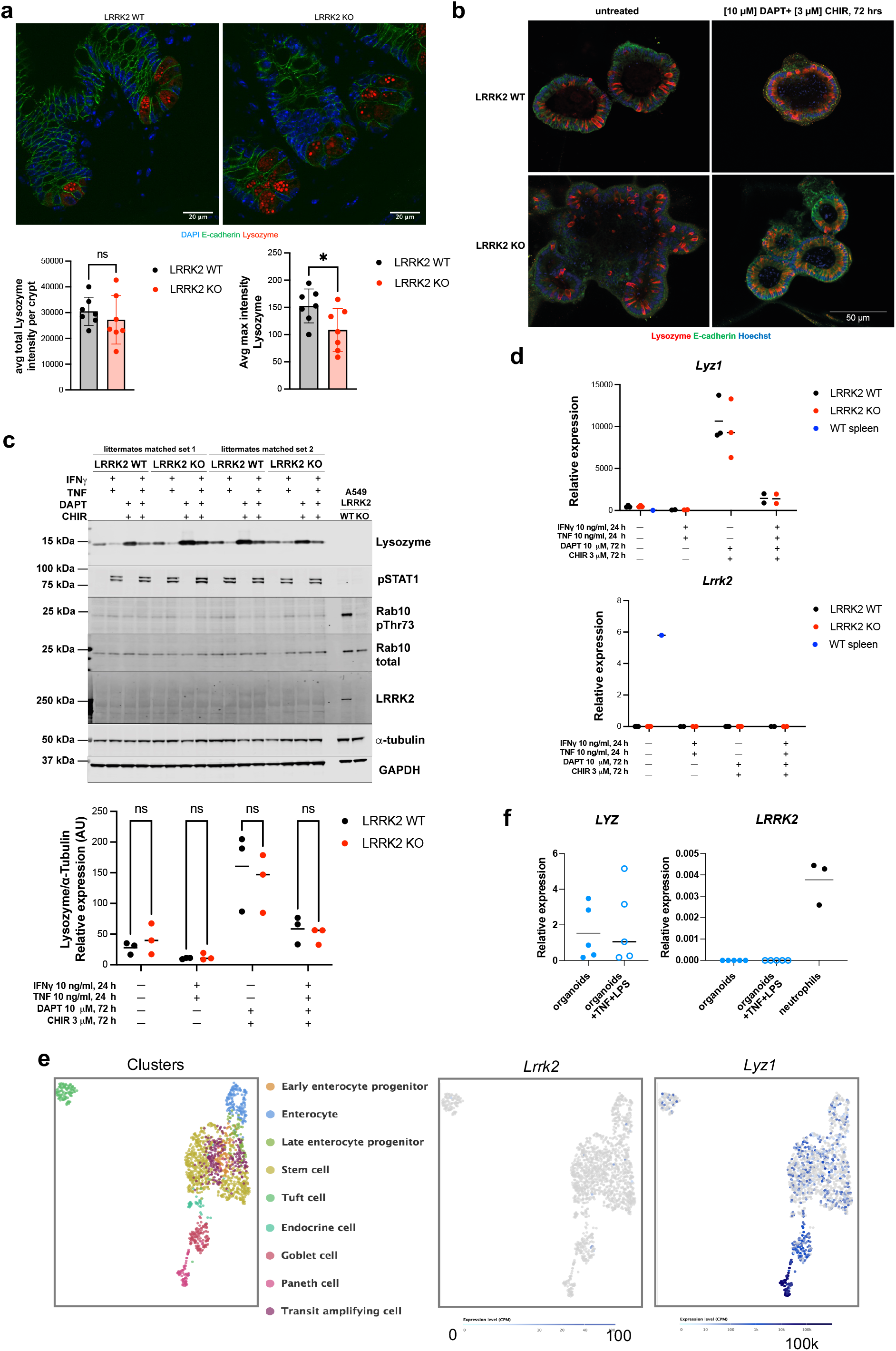
LRRK2 is not expressed, and is not required for lysozyme expression, in Paneth cells. **a)** Representative images of ileal sections from LRRK2 WT and LRRK2 KO mice stained as indicated. Quantification of total (left bottom) or maximal (right bottom) Lysozyme intensity per crypt from 8 random images per mouse in LRRK2 WT (n=7) and LRRK2 KO (n=7) mice. The experiment was reproduced four times with similar results. **b)** Representative images of mouse ileal organoids prepared from LRRK2 WT and LRRK2 KO mice, treated with a combination of DAPT (γ-secretase inhibitor) and CHIR 99021 (GSK-3 inhibitor) for 72h to enrich for Paneth cell differentiation, or left untreated, and stained as indicated. **c)** Littermate matched mouse ileum organoids from two sets of LRRK2 WT and LRRK2 KO were treated with or without the indicated inhibitors and/or cytokines and immunoblotted for LRRK2, Lysozyme, total and pT73 Rab10, pSTAT1 and the loading controls. Lysates from A549 LRRK2 WT and LRRK2 KO cell lines were used as controls for the antibodies against total and phosphorylated LRRK2 and Rab10. Quantitation of Lysozyme protein expression relative to tubulin from 3 biologically independent sets of organoids is shown below. **d)** mRNA expression of *lrrk2* and *lyz1* relative to housekeeping gene *tbp* was measured by qPCR in RNA extracted from organoids prepared and treated as in c). RNA from a LRRK2 WT spleen was used as a positive control for *Lrrk2* expression. **e)** Analysis of *Lrrk2* and *Lyz1* expression in murine intestinal epithelial cells in the Single Cell Portal from the Broad institute (data from Haber et al, 2017 (ref. 10)). **f)** Human ileal tissue-derived organoids (n=5 donors) were treated with or without TNF (100 ng/ml) and LPS (500 ng/ml) for 24 hrs. Relative expression of *LRRK2* and *LYZ* relative to housekeeping gene *B2M* was measured by qPCR. RNA from human peripheral blood neutrophils (n=3) was used as a positive control for *LRRK2* expression.

We next generated mouse ileal organoids from LRRK2 WT and KO littermate-matched mice to investigate the lysozyme expression specifically in the epithelial cells of the small intestine, in the absence of immune cells that express high level of LRRK2^7^. Organoids are formed from intestinal stem cells, and have all the epithelial lineages represented, including Paneth cells, but do not contain immune cells. Thus, organoids represent a valuable tool to study LRRK2 signaling exclusively in epithelial cells. We detected similar level of lysozyme staining in organoids grown from crypts extracted from both LRRK2 WT and LRRK2 KO mice (**Fig. 1b, left**). To further enrich for Paneth cell differentiation in organoids, we treated these ileal organoids with the γ-secretase inhibitor DAPT and the GSK-3 inhibitor CHIR 99021, which together drive stem cell differentiation to Paneth cells^8^. Even in these Paneth cell-enriched organoids, we found no impairment of lysozyme expression in crypts generated from LRRK2 KO mice, although there was a clear increase in lysozyme expression both at the protein and RNA level that confirmed the enrichment of Paneth cells (**Fig. 1b-d**). We could neither detect total LRRK2 protein nor the LRRK2-dependent phosphorylation of Rab10 at Thr73, using a highly selective and sensitive phospho-antibody^9^, in immunoblots (**Fig. 1c**) although we could detect LRRK2 in the corresponding WT splenic tissue from the same mice that were used to make the organoids (**Supplementary Fig. 1**). Expression of the *Lrrk2* gene is thought to be driven by IFNγ^10^. We therefore stimulated mouse ileal organoids with either IFNγ alone (not shown) or with IFNγ and TNF, another pro-inflammatory cytokine that is highly expressed in Crohn’s disease, but again we did not observe any signal for LRRK2 or any difference in lysozyme between the two genotypes (**Fig. 1c**). As expected, protein expression of lysozyme increased after enrichment of Paneth cells, and treatment with IFNγ caused a reduction in lysozyme protein in the organoids, consistent with IFNγ-induced lysozyme degranulation^11^. IFNγ stimulation was also confirmed by induction of phospho-STAT1. We further evaluated *Lrrk2* and *Lyz1* gene expression in the same set of LRRK2 WT and LRRK2 KO organoids: *Lrrk2* mRNA was not identified in organoids under any conditions, whereas it was measurable in splenocytes (**Fig. 1d**). *Lyz1* expression was detected and it increased in both genotypes equivalently when stimulated with DAPT and CHIR, and reduced when treated with IFNγ and TNF, as expected. LRRK2 protein was also not detected in deep proteomics dataset that identified >7700 proteins in sorted small intestinal epithelial cells from WT mice, although Paneth cell proteins such as lysozyme, defensins and Reg3γ were detected (**Supplementary table 1**). Further, we examined *Lrrk2* expression in published single cell RNA sequencing data from the murine small intestine^12^ on the Single Cell portal (https://singlecell.broadinstitute.org/single_cell), and found that *Lrrk2* was not detected in Paneth cells, although *Lyz1* expression was clearly identified and enriched in Paneth cells (**Fig. 1e**).

Separately, we analyzed *LRRK2* and lysozyme (*LYZ*) gene expression in human ileal tissue-derived organoids (n=5 individual donors), and in human neutrophils as a positive control for *LRRK2* expression^7^. Again, we could not detect *LRRK2* gene expression in the organoids without any stimulation, or with stimulation with TNF and LPS to mimic inflammatory conditions (**Fig. 1f**). *LYZ* was detected in the same conditions, indicating that Paneth cells were present in these organoids, but they did not express detectable levels of LRRK2.

Moreover, analysis of *LRRK2* expression in human intestinal cells^13^ in the Gut Cell Atlas from the Teichmann lab at the Wellcome Sanger Institute (https://www.gutcellatlas.org) revealed expression of *LRRK2* in stromal cells and immune cells, including B cells and myeloid cells, but not in any intestinal epithelial cell lineage. Thus, *LRRK2* is not expressed in Paneth cells in either the mouse or human gut.

The potential mechanism by which LRRK2 is thought to be involved in CD has been linked in multiple reviews^1,14,15^ to its putative role in regulating the sorting and the secretion of lysozyme in Paneth cells, as described in the study by Zhang *et al*^*4*^. Contrary to their study, we found no evidence that LRRK2 is expressed in Paneth cells. Moreover, LRRK2 does not directly regulate the expression or sorting of lysozyme in Paneth cells, as we see normal expression of lysozyme in LRRK2 KO Paneth cells. One conceivable explanation for the dissonance on lysozyme expression in LRRK2 KO Paneth cells with the previously published findings is the fact that production of the antibody that was used to stain for lysozyme expression by Zhang *et al*. (Abcam, ab36362), has been discontinued^16^ due to lack of specificity for lysozyme. We have also not been able to reproduce specific immunofluorescence staining of LRRK2 in intestinal tissue with the LRRK2 antibody (MJFF2, c41-2) used in their study. Thus, the use of these antibodies could have led to the contrasting conclusions in the study by Z. Liu and colleagues. We think it is extremely important to share our findings with the scientific community working towards a better understanding of the role of LRRK2 in CD where the molecular mechanisms involved are yet to be discovered, to broaden the scope of ongoing research beyond Paneth cells, perhaps to other immune cells in the gut that can also regulate Paneth cell biology.

## Acknowledgements

We acknowledge the support from Interline Therapeutics, and discussions with researchers at Interline, and at the University of Dundee.

## Author contributions

AT, DD, MoS performed experiments and analysed data, TH performed experiments, TM analysed data, MM provided reagents and human neutrophils and organoids and analysed data, AT and MS wrote the manuscript with input from all other authors.

## Data availability

All data presented in this study are shown in the figure. Raw data can be made available on request.

## Methods

### Ethics

Mice were bred and maintained with approval by the University of Dundee ethical review committee under a UK Home Office project license (PP2719506) in compliance with UK Home Office Animals (Scientific Procedures) Act 1986 guidelines.

Ethical approval for use of human ileal tissue was obtained from the Tayside Tissue Biorepository (Tissue Request No. 000657 via delegated authority (17/ES/0130) from The East of Scotland research ethics committee). All tissue was deidentified. Patients provided written consent. Full thickness segments of non-neoplastic terminal ileal mucosa from the resection margins were prepared by a pathologist from patients undergoing colorectal surgery for treatment of colorectal cancer. Ethical approval for the use of human blood derived neutrophils from healthy volunteers was obtained from the Schools of Medicine and Life Sciences Research Ethics Committee, University of Dundee (SMED REC 21/82).

### Mouse husbandry

Mice were maintained in a standard barrier facility on a 12hr light/dark cycle at 21°C in individually ventilated cages with sizzler-nest material and fed an R&M3 diet (Special Diet Services, UK) and filtered water ad libitum. Cages were changed at least every 2 weeks. *Lrrk2*^*-/-*^ (B6.129X1(FVB)-*Lrrk2*^*tm1*.*1Cai*^/J, LRRK2 KO) mice^6^ were obtained from Prof. Huaibin Cai (NIH, Bethesda), and backcrossed at least 10 generations in house to C57Bl/6J. Age-matched littermate controls were used for all experiments. Mice of both genders were used.

### Antibodies

The recombinant anti-lysozyme antibody [EPR2994(2)] (#ab108508), the recombinant phospho-Thr73-Rab10^9^ (#ab230261) and Rab10 total (#ab237703) were purchased from Abcam. The total Rab10 mouse antibody was from NanoTools (0680–100/ Rab10-605B11). Rabbit monoclonal antibody for LRRK2 phospho-Ser935 (#UDD2) was expressed and purified at University of Dundee as described previously^17^. The C-terminal total LRRK2 mouse monoclonal antibody was purchased from NeuroMab (clone N241A/34, #75-253). The rabbit monoclonal anti STAT1 phospho-Tyr701 antibody (clone 58D6, #9167) and the anti α-tubulin mouse monoclonal antibody (clone DM1A, mAb #3873) were purchased from Cell Signaling Technology. Anti-glyceraldehyde-3-phosphate dehydrogenase (GAPDH) antibody was from Santa Cruz Biotechnology (#sc-32233).

### Tissue immunofluorescence staining and microscopy of mouse ileum

Isolated mouse ileal tissues were fixed for 24h in 4% paraformaldehyde in PBS (pH = 7.4) at 4°C prior paraffin perfusion and embedding into paraffin blocks. 5 μm sections from paraffin-embedded tissues were deparaffinised using Histo-Clear xylene substitute, rehydrated and the antigen retrieval was performed in 10 mM Trisodium citrate containing 0.05% Tween 20, pH = 6.0, in a pressure cooker. The tissues were permeabilised for 20 min in 1% NP40 in PBS, blocked for 1 hour in PBS containing 2% Bovine Serum Albumin, 5% normal goat serum and 0.1% Triton X-100, and stained with the primary antibodies diluted in PBS containing 2% Bovine Serum Albumin and 0.1% TritonX100 (anti-lysozyme ab, 1:100; anti-E-cadherin ab, 1:150) at 4°C overnight, followed by staining with appropriate secondary antibodies (goat anti-rabbit IgG Alexa Fluor 568, Invitrogen #A-11036, and goat anti-mouse IgG Alexa Fluor Plus 488, Invitrogen #A32723), both highly cross-adsorbed to avoid any cross-species detection) diluted to 1:500 in PBS containing 2% Bovine Serum Albumin and 0.1% TritonX100 for 1 hour at room temperature. Washing was done with PBS throughout the procedure. The tissues were counterstained with 1 μg/ml DAPI and slides were mounted in Vectashield Vibrance antifade mounting media (Vector Laboratories, H-1700). Stained tissues were imaged on confocal Zeiss 710 microscope operated by ZEN software (Zeiss) using a 63x Plan Apochromat 1.4 NA oil objective, collecting a series of optical sections throughout tissue thickness. The maximal intensity projections from several optical sections were then generated in ImageJ. Alternatively, for quantitative analysis, 8 images were acquired on the same microscope with 20x Plan Apochromat 0.8 NA dry objective from at least 4 different tissue sections for each mouse. All images, including controls in which primary antibodies were omitted, were acquired using identical microscopy settings. Resulting three-channel images were further identically processed and assembled in OMERO.figure (https://www.openmicroscopy.org/omero/figure/).

### Image analysis

Three-channel confocal images were analysed in ImageJ/FIJI. The channel overlays were used to manually outline all visible crypts, and the total, mean and maximal fluorescence intensity in lysozyme channel was measured in each outlined crypt area. Eight images were analysed per mouse, with the total number of crypts between 85 and 112. The average values from all crypts of each mouse were then analysed to determine the mean total or maximal intensity per crypt per mouse. Statistical significance was calculated using t-test.

### Mouse ileal organoid culture, treatment and lysis for immunoblotting

Organoids were generated from the terminal ileum of mouse small intestine and established following the Stappenbeck method conditioned medium, produced in house^18^. Briefly, mice were culled by cervical dislocation (and confirmation by cut of femoral artery). The small intestine was dissected and flushed with cold DPBS (Gibco, #1419016) and cut longitudinally to expose the lumen. From about 2.5 cm above the caecum, the ileum was cut in small transverse sections and stored in 40 ml of complete RPMI medium (Gibco, #31870074) for crypt extraction. Working inside a bio-safety cabinet, the tissue sections were spun down at 1500x rpm for 5 min at room temperature and washed two times with DPBS. Tissue sections were incubated in 10 ml of DPBS containing 5 mM EDTA pH 8.0 at 4°C, in rotation, for 90 min. The cell suspension obtained was filtered away from the digested tissue using a cell strainer (100 μm). The tissues were transferred to a 15 ml tube and 10 ml of cold DPBS were added. Crypts were extracted mechanically by vigorously shaking the tube manually for 1 minute. The 15 ml tube was placed on ice to let the tissue settle down for 30 seconds. 6 ml of the supernatant containing the crypts was transferred to a new 15 ml tube. The volume was top up to 15 ml with cold DPBS and gently spun down at 100xg for 5 min. After aspiration of the supernatant, the crypts were gently resuspended in 5 ml of L-WRN medium (conditioned medium produced in house from L-WRN cells, ATCC, # CRL-3276). The suspension was spun at 100xg for 5 minutes, the medium aspirated and the crypts resuspended in 100 μl of L-WRN medium and directly added to 200 μl of Matrigel (Corning #356231) previously thawed on ice. Quickly, 30 μl drops of crypts:Matrigel were seeded in the middle of the wells of a 24-well plates, incubated 10 min inside an incubator at 37°C. After solidification of the Matrigel, 0.5 ml of warm L-WRN medium were added to each well. After 5-7 days, organoids were ready for subculturing and expansion in L-WRN medium.

At passage, 3 organoids were differentiated in Crypt-medium: Advanced DMEM/F12+++ medium (ADF, Gibco #12634-010) supplemented with L-glutamine and HEPES, B-27 (Gibco #17504-044)) and N-2 (Gibco #17502-048), 0.001 mM N-acetylcysteine, 50 ng/ml EGF, 100 ng/ml Noggin, 1 μg/ml R-spondin. DAPT (γ-secretase inhibitor, 10μM), CHIR99021 (GSK-3 inhibitor, 3μM). After 48h, TNF (10ng/ml) and IFNγ (10ng/ml)(both cytokines from Peprotech) were supplemented directly to the conditioned medium before being added to the wells. At the end of the time points, medium was aspirated and Matrigel scraped off as described above (inhibitors and cytokines were added at every step to avoid any possibility of wash out). Pellets were lysed in iced-cold lysis buffer (50 mM Tris–HCl, pH 7.4, 1% (by vol) Triton X-100, 10% (by vol) glycerol, 150 mM NaCl, 1 mM sodium orthovanadate, 50 mM sodium fluoride, 10 mM 2-glycerophosphate, 5 mM sodium pyrophosphate, 1 mg/ml microcystin-LR, and cOmplete EDTA-free protease inhibitor cocktail (Roche, #11836170001), snap frozen in liquid nitrogen and stored at –80°C until use. Lysates were thawed on ice for 30 minutes and clarified by centrifugation at 17000g for 20 min at + 4°C. Protein concentration was measured by Bradford assay (Bio-Rad Protein Assay Dye Reagent Concentrate #5000006) and samples prepared in 1X lithium dodecyl sulphate sample buffer (NuPage LDS Sample buffer 4× Invitrogen, #NP0008). To be able to load 25 to 35 μg of protein samples per lane in immunoblotting four wells per condition were pooled for lysis.

### Quantitative immunoblotting from mouse ileal organoids

25-35 μg of protein was loaded onto NuPAGE 4–12% Bis–Tris Midi Gel (Thermo Fisher Scientific, Cat# WG1403BOX) and electrophoresed with the NuPAGE MOPS SDS running buffer (Thermo Fisher Scientific, Cat# NP0001-02) at 200 V for 2 h. At the end of electrophoresis, proteins were transferred onto a nitrocellulose membrane (GE Healthcare, Amersham Protran Supported 0.45 mm NC) at 100V for 90 min on ice in transfer buffer (48mM Tris–HCl and 39mM glycine supplemented with 20% methanol). Transferred membrane was blocked with 5% (w/v) skim milk powder dissolved in TBS-T [20 mM Tris– HCl, pH 7.5, 150 mM NaCl and 0.1% (v/v) Tween 20] at room temperature for 1h. The membrane was typically cropped into three pieces, namely the ‘top piece’ (from the top of the membrane to 75 kDa), the ‘middle piece’ (between 75 and 30 kDa) and the ‘bottom piece’ (from 30 kDa to the bottom of the membrane). All primary antibodies were used at 1 μg/ml final concentration when the concentration was indicated, or diluted 1:1000, and incubated in TBS-T containing 5% (by mass) bovine serum albumin with exception of α-tubulin and GAPDH antibodies that were diluted 1:5000 and 1:2000, respectively. The top piece was incubated with the anti-STAT1-pTyr701 or the ] rabbit anti-pS935 LRRK2 antibody multiplexed with mouse anti-LRRK2 C-terminus total antibody. The middle piece was incubated with the anti-α-tubulin antibody. The bottom pieces were incubated with indicated anti-total and anti-pT73-Rab10 antibodies and anti-GAPDH antibody or anti-lysozyme antibody. Membranes were incubated in primary antibody overnight at 4°C. After three washes with TBS-T for 10 min each, membranes were incubated with secondary antibodies. The top and bottom pieces were incubated with goat anti-mouse IRDye 680LT (LI-COR, #926-68020) secondary antibody multiplexed with goat anti-rabbit IRDye 800CW (LI-COR, #926-32211) secondary antibody diluted in TBS-T (1:10000 dilution) for 1 h at room temperature, in the dark. The middle piece was incubated with goat anti-mouse IRDye 800CW (#926-32210) secondary antibody diluted in TBS-T (1:10000 dilution) at room temperature for 1 h, in the dark. Membranes were washed with TBS-T three times with a 10 min incubation for each wash and kept in the dark. Protein bands were acquired via near infrared fluorescent detection using the Odyssey CLx imaging system and quantified using the Image Studio software.

### Mouse ileal organoid immunofluorescence microscopy

For immunofluorescence, 15 μl Matrigel:mouse ileal organoids were seeded in μ-slides IbiTreat 8 well (#80826) and incubated in 200 μl of crypt medium. After 72 hrs incubation with [10 μM] DAPT and [3 μM] CHIR99021, the organoid domes were fixed: medium was aspirated, the domes were washed twice with 250μl DPBS and fixed with 200 μl of 4% PFA (by vol, final) for 30 min at 37ºC. To minimize the impact of PFA used for fixation on the Matrigel structure, the IbiTreat slides with the organoids and the PFA were incubated at 37ºC for 5 minutes to warm up. After 30 min, PFA was removed, the domes were washed once with DPBS and permeabilized with 1% (by vol) Triton X-100 in DPBS at RT. The slides were washed once with blocking buffer (1% (by mass) BSA, 0.3% normal goat serum (by vol), 0.2% (by vol) Triton X-100 in DPBS) and blocked in blocking buffer for 2 hrs. Primary antibodies were diluted in working buffer (0.1% (by mass) BSA, 0.3% (by vol) normal goat serum, 0.2% (by vol) Triton X-100 in DPBS) and incubated over night at RT. The domes were washed 5 times with 200 μl of working buffer and incubated overnight, in the dark, with secondary antibodies diluted 1:500 in working buffer. The next day, after 5 washes with 200 μl of working buffer, the domes were incubated for 20 min at RT in the dark with Hoechst 33342 (Invitrogen, #H3570). The domes were washed again 5 times with 200 μl of working buffer and prepared for imaging with 200 μl of ProLong Glass mounting medium (Invitrogen, #P36984) and let set overnight at RT in the dark. Organoids were stained with the anti-lysozyme antibody for imaging diluted 1:1000 (Dako, #A0099) and visualized with goat antirabbit Alexa fluor 568 (Invitrogen #A-11036), the anti-E-cadherin antibody (BD Biosciences, #610182) was diluted 1:50 and then visualized with goat anti-mouse Alexa fluor 488 (Invitrogen #A32723). The images were collected on an LSM710 laser scanning confocal microscope (Carl Zeiss) using the 20X Plan Apochromat 0.8 NA objective.

### mRNA extraction from murine organoids

A minimum of four wells of mouse organoids were pooled and lysed following the protocol of the PureLink RNA Mini Kit (Ambion, Life Technologies, #12183025). Briefly, after 2 washes in cold DPBS, organoids were resuspended in 15 ml of cold DPBS and clarified from residual Matrigel at 500xg for 5 min at 4ºC. The supernatant was carefully aspirated and lysed in 600 μl of the lysis buffer supplemented with β-mercaptoethanol (10 μl (by vol) in 10 ml of Lysis buffer). mRNA was eluted in 30 μl of RNase-free water from the kit.

50 mg of mouse spleens were homogenized in 500 μl of the same lysis buffer using the TissueLyser LT (Qiagen, #85600) (50 sec for 2 min) and filtered through an 18-gauge needle. mRNA was eluted in 50 μl of RNase-free water.

### Retrotranscription and Real Time PCR in mouse derived tissue

cDNA synthesis and Real-Time PCR were performed following the manufacture instructions of the PrimeScript RT reagent Kit with gDNA Eraser kit (Takara, #RR047A). For the Real-Time PCR reaction, the cDNA obtained was diluted 1:5 in nuclease-free water, mixed to 10 μM Forward primer, 10 μM Reverse primer and TB Green Premix Ex Taq II (Tli RNaseH Plus) (Takara, #RR820W). Real-Time PCR was set up in a 384-well plate and run in a CFX384 Real-Time PCR System (BioRad) using the following protocol: initial denaturation at 95ºC for 30 sec, 40 cycles of PCR at 95ºC for 5 sec - 60ºC for 1 min followed by 5 min at 65ºC and 5 min at 95ºC. The sequences of the primers used for Real-Time PCR are the following and were purchased from Eurofins: *Lrrk2* Forward 5’-cagcttcagaagggacaagg – 3’ and Reverse 5’-aaggctgcgttctcaggata – 3’, *Lyz1* Forward *5’-* GAGACCGAAGCACCGACTATG - 3’, Reverse 5’-CGGTTTTGACATTGTGTTCGC-3’, *tbp* Forward 5’-GGGGAGCTGTGATGTGAAGT – 3’ , Reverse 5’ -CCAGGAAATAATTCTGGCTCAT – 3’. Gene expression relative to the housekeeping gene was calculated using the 2^-ΔCt^ method.

### Sorting of mouse intestinal epithelial cells for proteomics

For proteomics experiments, 3 biological samples from 3 mice (male, C57BL/6J, aged 10-12 weeks) were generated. Intestines were dissected from proximal duodenum to terminal ileum and flushed with HBSS (Ca^2+^ and Mg^2+^ free) to remove luminal contents. Intestines were cut longitudinally, Peyer’s patches removed, washed vigorously twice in HBSS and then cut into 5-10mm pieces. The pieces were incubated in HBSS with 2% FCS, 5mM DTT, 5mM EDTA, 1 μM Y-27632 dihydrochloride (ROCK inhibitor, Tocris, #1254) on a horizontal shaker for 30 minutes at 37ºC. Pieces were spun down and vortexed in Advanced DMEM/F12+++ (ADF, see above) for 3min, and the released cells filtered through a 100μm filter. Cells where there treated with Gentle cell dissociation reagent (Stemcell Technologies, #100-0485) +1 μM Y-27632 for 10 minutes at RT. After washing, cells were stained with Live/Dead dye Aqua, CD45 APC (1:200), EPCAM-PE (1:800) in ADF containing 10 μM Y-27632 and DNase. Cells were sorted according to their forward scatter and side scatter, dead cells excluded, and EPCAM-hi, CD45-negative cells were selected as epithelial cells.

### Sample preparation for mass spectrometry

Epithelial cell pellets containing 1 million cells were lysed and prepared as described^19^ with the following changes: Pellets were lysed in 400μL lysis buffer (4% SDS, 10mM TCEP, 50mM TEAB (pH 8.5)). After peptide clean-up according to the SP3 protocol, samples were resuspended in 2% DMSO and 5% formic acid and fractionated by off-line high pH (9.5) reverse phase chromatography into 16 concatenated fractions. Samples were analysed by mass spectrometry at the FingerPrints facility, University of Dundee, where each fraction was analysed by label-free quantification (LFQ) using an LTQ-Orbitrap Velos Pro mass spectrometer running Xcalibur software (Thermo Scientific) with a 240-minute gradient per fraction.

### Processing and statistical analysis of proteomics data

The MS raw data files were processed with MaxQuant version 1.6.8.0 and Perseus v1.5.2.6, against mouse reviewed proteome from Swissprot with isoforms, downloaded in Aug 2019. Minimum peptide length was set to 6, and proteins were quantified on unique+razor peptides with the following modifications included : Oxidation (M); Acetyl (Protein N-term); Deamidation (NQ). The data set was filtered to remove proteins categorised as “contaminants” and “reverse”. Analysed data is presented in Supplementary table 1.

### Human ileal tissue derived organoid culture

A human ileum-derived epithelial organoid model was established, optimised from our previously published human colon organoid method^20,21^ based on a published protocol^22^. In summary, human ileal mucosa was incubated in 5mM EDTA buffer on ice for 5 min with gentle agitation, shaken for 5 minutes to dislodge the crypts, then passed through a 100 μM cell strainer. Following centrifugation and washing, crypts were suspended in Matrigel, supplemented with 1:100 Jagged-1-peptide (Cambridge Bioscience, #ANA61298) at 200 crypts/15 μL Matrigel. A total of 15 μL crypt Matrigel mixture per well was dispensed into a pre-warmed delta-surface 24-well plate, and the Matrigel was polymerised by inverting plate at 37 °C for 15 min. For initiating, maintaining and differentiation of human ileal organoid culture, a 50% conditioned media (1:1 dilution in advanced D-MEM/F12 base media) was generated from genetically modified L-WRN mouse fibroblast cell line (ATCC, #CLR-3276™, ATCC, LGC Standards, Middlesex, UK) cultured in advanced D-MEM/F12 (Thermofisher, #12634010) with 10% FCS. L-WRN cells were selected for 24 hrs by incubation with 0.5 mg/mL hygromycin-B (Thermofisher, #10687010) and 0.5 mg/mL G-418 Sulfate (Thermofisher, #10131035). Initiation media contained 1% bovine serum albumin (Thermofisher™, B14) , 2 mM Glutamax (Gibco™, 35050061), 10 mM HEPES (Gibco™,#15630080), 1× N2 Supplement (Gibco™, #17502048), 1× B27™ Supplement (Gibco™, #17504044), 1 mM N-acetylcysteine (Sigma-Aldrich , #A9165-5G), 50 ng/mL rhEGF (Peprotech, #AF-100-15-500), 10 mM nicotinamide, 500 nM A83-01 (Tocris, #2939), 10 nM, prostaglandin E2 (Tocris, 2296), 10 nM [Leu-15]-gastrin1 (Sigma-Aldrich, G9145), 10 μM SB202190 (Sigma-Aldrich, #S7067), 2.5 μM Thiazovinin (Stemcell Technologies, #72252), 10 μM Y-27632 dihydrochloride (Tocris, #1254), 2.5 μM CHIR99021 (Tocris, #4423), and 100 μg/mL Primocin® (InvivoGen, #ant-pm-05). Maintenance media was prepared with the same constituents minus 10 μM Y-27632 dihydrochloride and 2.5 μM CHIR99021. Differentiation media was prepared as per initiation media minus 10 mM nicotinamide, 10 μM SB202190, 10 μM Y-27632 dihydrochloride and 2.5 μM CHIR99021. Prior to experiments, organoids were cultured in 100% DMEM/F12 with the same additives as per differentiation media, minus 10 nM prostaglandin E2 and 2.5 μM Thiazovinin and with added human recombinant Wnt3A 50 ng/mL (R&D Systems, #5036-WN-010/CF), R-Spondin-1 250 ng/mL (Peprotech, #120-38) and recombinant human noggin 50 ng/mL (Peprotech, #120-10C). Organoids were cultured at 37 °C in 5% CO2 in initiation media for 72 hr, then replaced with maintenance media, refreshed every 2-3 days. Before experimental stimulations, organoids were plated on delta-surface 96-well plates and cultured in initiation media for 2 days, maintenance media for 3 days, differentiation media for 4 days until maturity before swapping to fully recombinant differentiation media for 24 hrs before experimental use. To simulate inflammatory conditions, organoids were stimulated with TNF (100 ng/ml, Peprotech, #300-01A) and LPS (100 ng/ml) for 24 hrs.

### Human neutrophil isolation

Human neutrophils were extracted by negative selection from the blood of healthy volunteers using MACSxpress® Whole Blood Neutrophil Isolation Kit (Miltenyi, #130-104-434) and MACSxpress separator (Miltenyi, #130-098-308), incorporating the MACSxpress erythrocyte depletion kit (Miltenyi, 130-098-196). Human neutrophils were cultured in RPMI-1640 media (Gibco, #52400017) + 10% FCS + 1% penicillin/streptomycin and the purity of isolated neutrophils was confirmed as >98% CD45+ CD15+ via flow cytometry (anti-CD45-FITC, 555482 and anti-CD15-PE, 555402, BD Biosciences).

### Gene expression analysis in human ileal organoids and neutrophils

RNA was extracted from human tissue derived organoids and neutrophils using the RNeasy® Plus Mini Kit (Qiagen, #74104) with cDNA prepared using a SuperScript™ VILO™ cDNA synthesis kit (Invitrogen, #11756050). RT-qPCR using TaqMan assays (Applied Biosystems, #4351372) and TaqMan™ Fast Advanced Master Mix (Applied Biosytems, #4444556) characterised *LRRK2* (Hs01115057_m1) and *LYZ* (Hs00968202_m1) gene expression normalised to *B2M* (Hs00187842_m1) on a StepOnePlus™ real-time PCR system (Applied Biosystems, Thermo Fisher Scientific UK Ltd., Loughborough, UK). Gene expression relative to the housekeeping gene *B2M* was calculated using the 2^-ΔCt^ method.

---

**Supplementary Figure 1.**
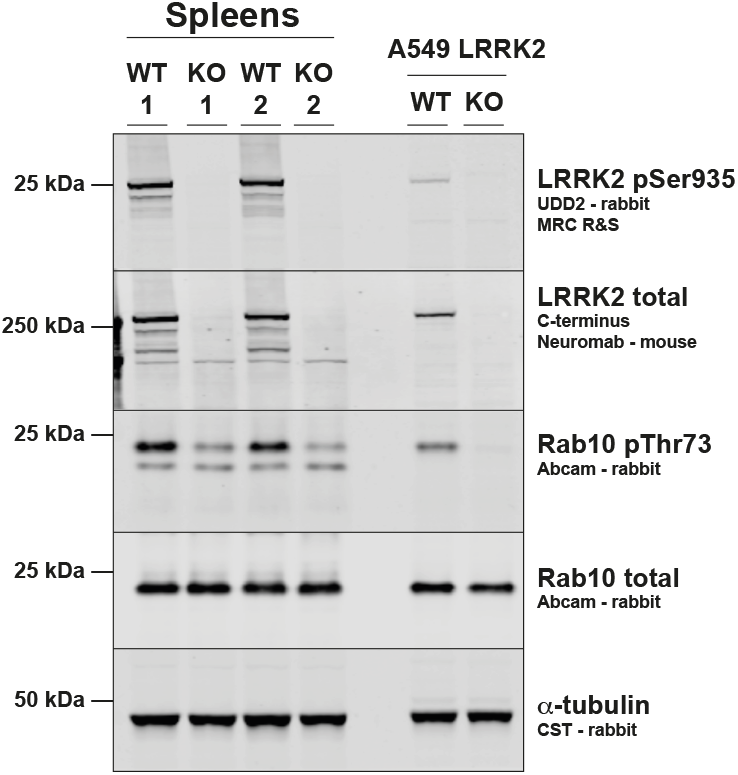
Control immunoblot to show the deletion of LRRK2 in the *Lrrk2*^*-/-*^ mice used to generate ileal organoids shown in Figure 1. The spleens were dissected and snap frozen in liquid nitrogen and stored at –80°C until use. 30 mg of each spleen were homogenized using a tissue pulveriser (cellcrusher-mini, https://cellcrusher.com/mini-tissue-pulverizer-3/). The homogenates were left on ice for 30 minutes and clarified by centrifugation at 17000x g for 30 min at 4°C. Protein concentration was measured by Bradford assay (Bio-Rad Protein Assay Dye Reagent Concentrate #5000006) and samples prepared for immunoblotting as above. 40 μg of samples were loaded.

